# Protection of Hamsters Challenged with SARS-CoV-2 Variants of Concern by Two Doses of MVC-COV1901 Vaccine Followed by a Single Dose of Beta Variant Version of MVC-COV1901

**DOI:** 10.1101/2021.09.29.462344

**Authors:** Tsun-Yung Kuo, Chia-En Lien, Yi-Jiun Lin, Meei-Yun Lin, Chung-Chin Wu, Wei-Hsuan Tang, John D. Campbell, Paula Traquina, Ya-Shan Chuang, Luke Tzu-Chi Liu, Jinyi Cheng, Charles Chen

## Abstract

The current fight against COVID-19 is compounded by the Variants of Concern (VoCs), which can diminish the effectiveness of vaccines and potentially increase viral transmission and severity of disease. MVC-COV1901 is a protein subunit vaccine based on the prefusion SARS-CoV-2 spike protein (S-2P) and is adjuvanted with CpG 1018 and aluminum hydroxide. In this study, we used the Delta variant to challenge hamsters inoculated with S-2P from the Wuhan wildtype and the Beta variant in two-dose or three-dose regimens. Two doses of wildtype S-2P followed by the third dose of Beta variant was shown to induce the highest neutralizing antibody titer against live SARS-CoV-2 of the wildtype and all current VoCs, as well as improved neutralization against Omicron variant pseudovirus compared to three doses of wildtype S-P. All regimens of vaccination were able to protect hamsters from SARS-CoV-2 Delta variant challenge and resulted in reduced lung live virus titer and pathology. Three doses of vaccination also significantly reduced lung viral RNA titer, regardless of whether the wildtype or Beta variant S-2P was used as the third dose. Based on the immunogenicity and viral challenge data, two doses of wildtype S-2P followed by the third dose of Beta variant S-2P induced potent antibody immune responses against the VoCs.

## Introduction

As of 2022, the COVID-19 pandemic shows no sign of abating despite the fact that over five billion doses of vaccines have been administered worldwide. The emergence of variants has undoubtedly played an important role in facilitating the global spread of COVID-19 as recent global dominance of Delta variant followed by Omicron variant have shown [1]. The WHO currently lists five VoCs: Alpha (B.1.1.7), Beta (B.1.351), Gamma (P.1), Delta (B.1.617.2), and Omicron (B.1.1.529), and two Variants of Interest (VoIs): Lambda (C.37) and Mu (B.1.621) [2]. In addition to increased transmission, the VoCs are known to reduce the *in vitro* neutralizing capability of currently available vaccines through mutations on the spike protein, especially in the receptor binding domain (RBD), and the most recent VoC Omicron has shown to be highly refractory to neutralization by convalescent and vaccine-induced antibodies [3–6]. These *in vitro* data have been shown to have a tangible impact on public health with reports of diminished vaccine efficacy, particularly among those infected with the Beta, Delta, and Omicron variants [5–10]. Instead of developing variant-based vaccine or a universal SARS-COV-2 vaccine, administration of booster vaccination has shown to be effective against the VoCs [11–14]

Medigen’s MVC-COV1901 is a subunit vaccine based on a stabilized prefusion S-2P protein adjuvanted with CpG 1018 and aluminum hydroxide [15]. This vaccine has been shown to be safe and highly immunogenic in both hamster challenge studies and clinical trials [16–18], and has been approved for emergency use in Taiwan. The vaccine is given intramuscularly as two doses separated by four weeks [19]. We have previously shown that two doses induce neutralizing antibodies against SARS-CoV-2 variants with a tendency of higher immunogenicity at higher dose levels [20]. We have also found that a third dose of this vaccine administered to rats increased neutralizing antibody titers against the Beta variant compared to just two doses [20]. The current study expands on our previous findings to investigate the immunogenicity of third dose booster against current VoCs.

## Methods

### Animals and ethical statements

Female golden Syrian hamsters aged 8-10 weeks at study initiation were obtained from the National Laboratory Animal Center (Taipei, Taiwan). Animal immunizations were conducted in the Testing Facility for Biological Safety, TFBS Bioscience Inc., Taiwan. At three weeks following the final immunization, the animals were transferred to Academia Sinica, Taiwan, for SARS-CoV-2 challenge. All procedures in this study involving animals were conducted in a manner to avoid or minimize discomfort, distress, or pain to the animals and were carried out in compliance with the ARRIVE guidelines (https://arriveguidelines.org/). All animal work in the current study was reviewed and approved by the Institutional Animal Care and Use Committee (IACUC) with animal study protocol approval number TFBS2020-019 and Academia Sinica (approval number: 20-06-1483).

### Immunization and challenge of hamsters

The study design is outlined in Figure 1. The hamsters were split into the following six groups with n = 10 for each group (Table 1):

**Figure 1.**
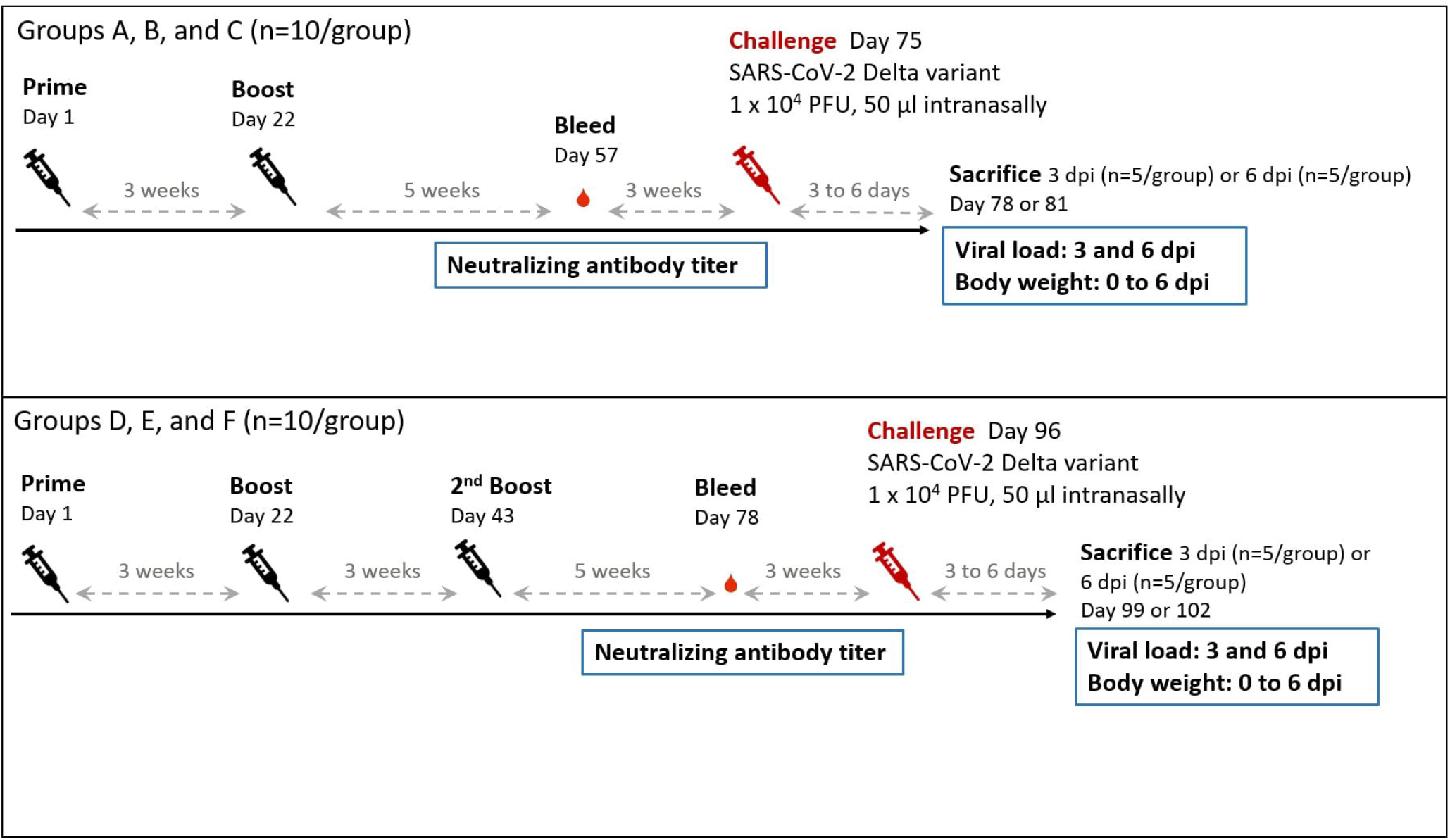
Study design of the hamster challenge study. Hamsters (N=10 per group) were immunized twice (groups A, B, and C) or three times (groups D, E, and F) at three weeks apart and serum samples were taken for immunogenicity assays five weeks after the final immunization. Eight weeks after the final immunization, hamsters were challenged with 10^4^ PFU of SARS-CoV-2 Delta variant. The animals were euthanized on the third or sixth day after infection for necropsy and tissue sampling to determine viral load. Body weight of individual hamster were tracked daily up to the time of sacrifice.

**Table 1.**
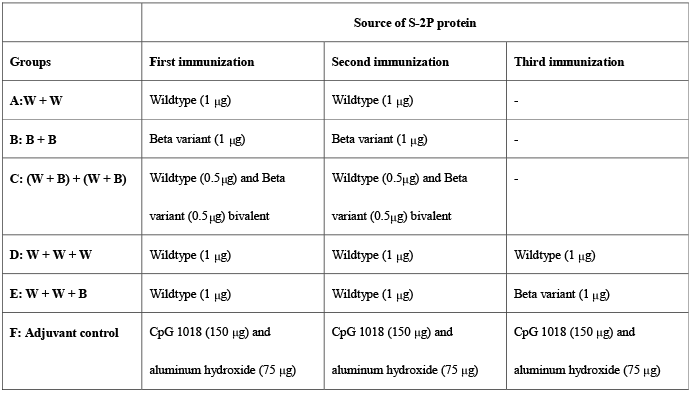
Hamster group and S-2P dosing.

| Groups | Source of S-2P protein |  |  |
| --- | --- | --- | --- |
|  | First immunization | Second immunization | Third immunization |
| **A: W + W** | Wildtype (1 µg) | Wildtype (1 µg) | - |
| **B: B + B** | Beta variant (1 µg) | Beta variant (1 µg) | - |
| **C: (W + B) + (W + B)** | Wildtype (0.5µg) and Beta variant (0.5µg) bivalent | Wildtype (0.5µg) and Beta variant (0.5µg) bivalent | - |
| **D: W + W + W** | Wildtype (1 µg) | Wildtype (1 µg) | Wildtype (1 µg) |
| **E: W + W + B** | Wildtype (1 µg) | Wildtype (1 µg) | Beta variant (1 µg) |
| **F: Adjuvant control** | CpG 1018 (150 µg) and aluminum hydroxide (75 µg) | CpG 1018 (150 µg) and aluminum hydroxide (75 µg) | CpG 1018 (150 µg) and aluminum hydroxide (75 µg) |

**Table 2.** List of mutations for spike sequences used in construction of variant pseudoviruses. Sequences were chosen from the most commonly found mutation in circulation according to Coronavirus3D at the time of construction (https://coronavirus3d.org/).

| WHO designation | PANGO lineage | Mutations |
| --- | --- | --- |
| Alpha | B.1.1.7 | L18F, del69, del144, N501Y, D614G, P681H |
| Beta | B.1.351 | L18F, del242, K417N, E484K, N501Y, D614G, 701V |
| Delta | B.1.617.2 | T95I, L452R, T478K, D614G, P681R |
| Omicron | B.1.1.529 | A67V, del69-70, T95I, G142D, del143-145, del211, L212I, ins214EPE, G339D, S371L, S373P, S375F, S477N, T478K, E484A, Q493R, G496S, Q498R, N501Y, Y505H, T547K, D614G, H655Y, N679K, P681H, D796Y, N856K, Q954H, N969K, L981F |

Vaccination was administered to hamsters via intramuscular injection in quadriceps femoris muscle of left and right legs (50 μL each leg for a total of 100 μL per dose). Hamsters in group A were vaccinated on days 22 and 43 with 1 μg of S-2P protein derived from the wildtype. Hamsters in group B were vaccinated on days 22 and 43 with 1 μg of S-2P protein derived from Beta variant. Hamsters in group C were vaccinated on days 22 and 43 with a mixture of the wildtype (0.5 μg) and Beta variant (0.5 μg) of S-2P protein (bivalent vaccine). Hamsters in group D were vaccinated on days 1, 22, and 43 with 1 μg of S-2P protein derived from the wildtype. Hamsters in group E were vaccinated on days 1 and 22 with 1 μg of wildtype S-2P protein, and on day 43 with 1 μg of S-2P protein derived from the Beta variant. Hamsters in group F served as an adjuvant control and were vaccinated with only 150 μg of CpG 1018 and 75 μg of aluminum hydroxide (alum) on days 1, 22 and 43. All immunization with S-2P were adjuvanted with 150 μg of CpG 1018 and 75 μg of alum.

Serum samples were collected five weeks after the final immunization and immunogenicity was determined by neutralization assay with SARS-CoV-2 virus and the variants. Approximately three weeks after the serum sampling (53 days after the final immunization), hamsters were challenged with the SARS-CoV-2 Delta variant (TCDC#1144) and then sacrificed at 3 d.p.i. (n = 5 per group) or 6 d.p.i. (n = 5 per group) for analyses of lung viral loads, lung TCID_50_. Body weight of individual hamsters were tracked daily up to the time of sacrifice. Necropsy were performed with lungs of euthanized hamster and histopathology sectioning, staining, and scoring were done as described previously [13]. The left lung of hamsters was isolated and fixed in 4% paraformaldehyde for one week. After fixation the lung was trimmed, processed, embedded, sectioned, and stained with Hematoxylin and Eosin (H&E), followed by microscopic examination. The lung section was evaluated with a lung histopathological scoring system in which a micrograph of a section was divided into nine equal squares and a score was assigned from 0 (normal) to 5 (most severe histopathology) for each of the nine squares and the average of scores from the squares was calculated to represent the score of the animal [13].

### Live SARS□CoV□2 and pseudovirus neutralization assay

SARS-CoV-2 virus strains including Wuhan wildtype (hCoV□19/Taiwan/4/2020, GISAD EPI_ISL_411927), Alpha (B.1.1.7, hCoV□19/Taiwan/792, GISAD EPI_ISL_1381386), Beta (B.1.351, hCoV□19/Taiwan/1013), Gamma (P.1, hCoV□19/Taiwan/906), and Delta (B.1.617.2, hCoV□19/Taiwan/1144) variants were used in live virus neutralization assay as described previously [17]. Briefly, two-fold serial dilutions of serum samples were mixed with equal volume of 100 TCID_50_/50 μL of virus and incubated at 37 °C for one hour. The mixture was added to the cell plates containing the Vero E6 cells and incubated at 37 °C, for four to five days. The neutralising titre (NT50) was defined as the reciprocal of the highest dilution capable of inhibiting 50% of the cytopathic effect (CPE).

Pseudovirus with lentivirus pseudotyped with S proteins of the wildtype, Alpha, Beta, Delta, and Omicron variants were conducted as previously described [12]. Two-fold serial dilution of serum samples were mixed with equal volume of pseudovirus (1000 transduction unit) and incubated before adding to the plates with HEK293-hAce2 cells. The amount of pseudovirus entering the cells was expressed relative luciferase units (RLU). Fifty percent inhibition dilution titers (ID_50_) were calculated with uninfected cells as 100% neutralization and cells transduced with virus as 0% neutralization. A list of mutations in the spike sequences used in the construction of pseudoviruses is included below.

### Viral RNA quantification and cell culture infectious assay (TCID_50_)

Quantification of lung viral load by real-time PCR and TCID_50_ assay were performed as previously reported [13]. Briefly, RNA from lung tissue was extracted using RNeasy Mini Kit (QIAGEN Germany). RNA sample was reverse transcribed using the Superscript III one-step RT-PCR system with Platinum Taq Polymerase (Thermo Fisher Scientific, USA) and a set of primers targeting the SARS-CoV-2 envelope (E) gene was used to amplify by TaqMan real-time RT-PCR.

TCID50 was determined from homogenized tissue of middle, inferior, and post-caval lung lobes of hamsters. Ten-fold serial dilutions of sample were added onto Vero E6 cell in quadruplicate and incubated for 4 days at 37°C. Cells were fixed with 4% formaldehyde and stained with 0.5% crystal violet for visualization.

### Statistical analysis

The comparisons between neutralizing antibody titers were performed using Kruskall-Wallis test with corrected Dunn’s multiple comparisons test, two-way ANOVA with Dunnett multiple comparison test, and unpaired Mann-Whitney U test. Correlation between SARS-CoV-2 viral genome copy numbers and NT50 titers against the Delta variant was estimated with Spearman’s rank coefficient and simple linear regression. Statistical analysis was conducted with Prism 6.01 (GraphPad).. * = p < 0.05, ** = p < 0.01, *** = p < 0.001, **** = p < 0.0001

## Results

### Induction of neutralizing antibodies against VoCs in hamsters immunized with two doses of S-2P derived from original MVC-COV1901, Beta variant, or a bivalent combination of both

We have previously shown that there was approximately a 7-fold reduction in the neutralizing antibody titer against the Beta variant in the sera of individuals who received two doses of MVC-COV1901 during a phase I trial [20]. We subsequently developed a Beta variant version of S-2P produced by a stable CHO cell clone and adjuvanted with 750 μg of CpG 1018 and 375 μg of aluminum hydroxide. We have previously established that two doses using one-fifth the amount of either low dose or high dose of MVC-COV1901 were sufficient to protect hamsters from SARS-CoV-2 infection [16]. Since the Delta variant has become prevalent worldwide, we investigated the protective effect of MVC-COV1901 derived from Wuhan wildtype (W), its Beta variant version (B) of S-2P, and the wildtype/Beta S-2P bivalent vaccine in hamsters challenged with the Delta variant. We first examined the neutralizing antibody titers from hamsters immunized with two doses of 1 μg wildtype S-2P adjuvanted with 150 μg CpG 1018 and 75 μg aluminum hydroxide (Group A shown as W + W). As shown in Figure 2, at five weeks after the second injection, Group A hamsters showed a reciprocal 50% neutralizing antibody titer (NT_50_) GMT of 2201, 581, 166, 193, and 742 against the wildtype, Alpha, Beta, Gamma, and Delta variants, respectively. Compared to the neutralizing titer against the wildtype, those against the Alpha, Beta, Gamma, and Delta variants showed a 3.79-, 13.30-, 11.39-, and 2.97-fold reduction, respectively. This demonstrated that two doses of S-2P derived from wildtype was relatively effective against the Alpha and Delta variants. However, the effectiveness was significantly reduced against the Beta and Gamma variants.

**Figure 2.**
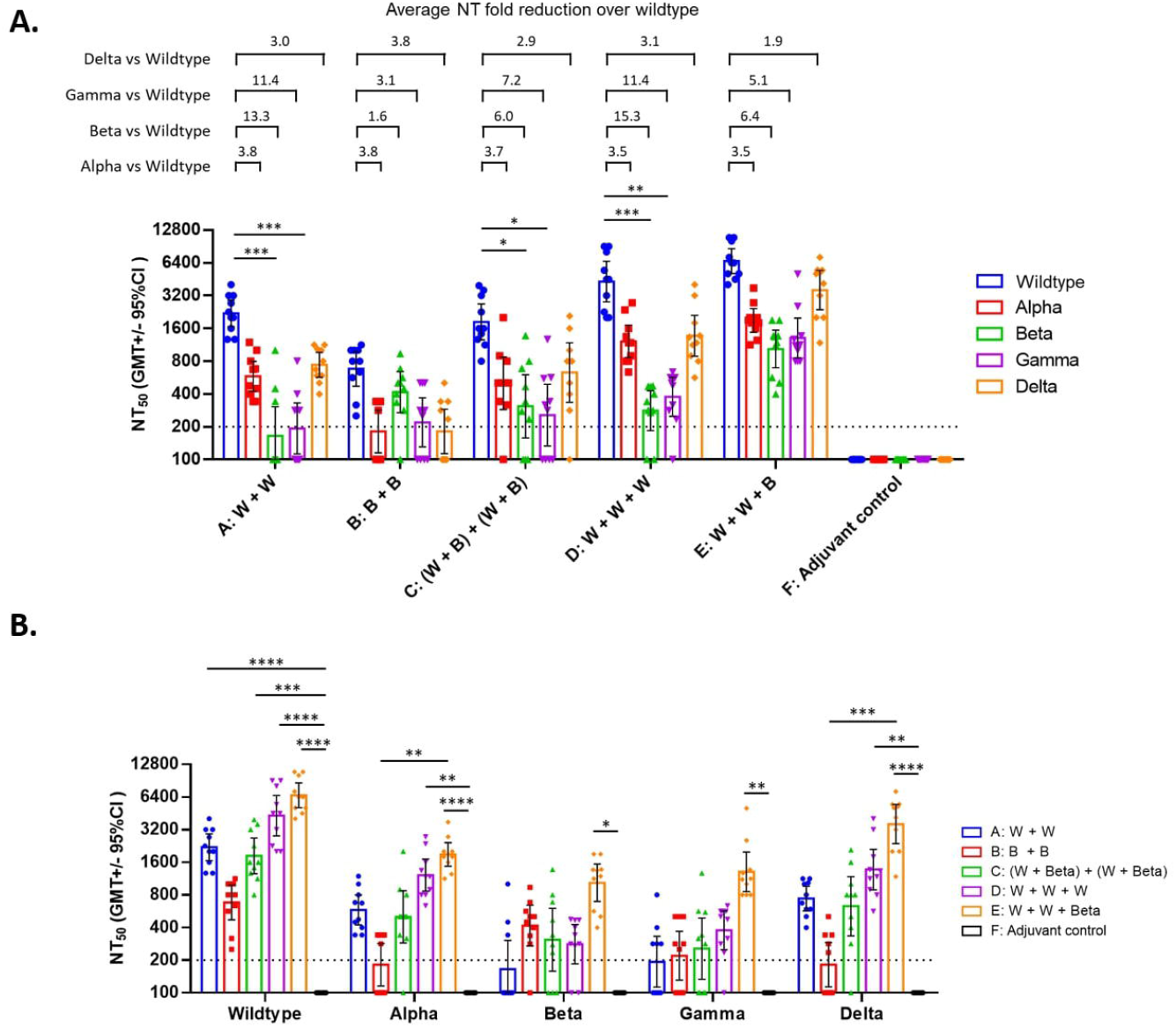
Neutralizing antibody titers with live SARS-CoV-2 neutralization assay in hamsters five weeks after the final immunization. Hamsters were immunized as in Figure 1. Five weeks after the final immunization (second immunization for groups A, B, and C; third immunization for groups D, E, and F), serum samples were taken for neutralization assays against live SARS-CoV-2 Wuhan wildtype strain and Alpha, Beta, Gamma, and Delta variants. The results were **A:** Grouped according to vaccine regimen; **B:** Grouped according to variants. Bars indicate NT_50_ GMT with individual values displayed as symbols and error bars showing the 95% confidence intervals. Average fold reduction in GMT of variants against the wildtype strain were calculated and shown above brackets above the corresponding bars. W: Wildtype S-2P; B: Beta variant S-2P; W + B: bivalent mixture of widltype and Beta variant S-2Ps. Dotted line indicate the starting dilution (200) and all values below 200 are tabulated as 100. Statistical significance was calculated with Kruskal-Wallis test with corrected Dunn’s multiple comparisons test. * = p < 0.05, ** = p < 0.01, *** = p < 0.001, **** = p < 0.0001

At the same time, we examined the neutralizing antibody titers from hamsters immunized with two doses of 1 μg of the Beta variant version of S-2P combined with 150 μg CpG 1018 and 75 μg aluminum hydroxide - Group B (B + B). Figure 2 shows that two doses of the adjuvanted Beta variant S-2P induced a satisfactory NT_50_ GMT of 681 and 417 against the wildtype and Beta variant, respectively. However, the neutralizing titers of this regimen was less than desirable against the Alpha, Gamma, and Delta variants that were 181, 219 and 182, respectively.

We also explored the neutralizing antibody responses of bivalent vaccine (wildtype + Beta variant) in Group C hamsters [shown as (W + B) + (W + B)]. The bivalent vaccine induced a similar degree of neutralizing antibody titers against the wildtype, Alpha, and Delta variants to that of the W+W group. This combination fared better against the Beta and Gamma variants than that of the W+W group; however, neutralization titer against these variants in several individual hamsters were less than 200.

### Induction of neutralizing antibodies against VoCs in hamsters immunized with two doses of MVC-COV19 combined with a third dose of MVC-COV1901 or its Beta variant version of S-2P

We previously found that neutralizing antibody titers against the Beta variant increased substantially in rats that had received three rather than two doses of MVC-COV1901 [20]. We therefore immunized hamsters with a third dose of one-fifth amount of MVC-COV1901, referred to as Group D (W + W + W), and examined the antibody responses against VoCs. As shown in Figure 2, five weeks after the third dose in Group D hamsters, NT_50_ GMTs were 4302, 1217, 281, 377, and 1368 against the wildtype, Alpha, Beta, Gamma, and Delta variants, respectively. The neutralizing titers against the Alpha, Beta, Gamma, and Delta variants had a 3.54-, 15.30-, 11.41- and 3.14-fold decrease, respectively, compared to that of the wildtype. Compared to Group A hamsters (which only received two doses), the neutralizing antibody titers in Group D hamsters against VoCs increased substantially with the additional third dose. The third dose not only increased the neutralizing antibody titers against the Delta variant but also boosted neutralizing antibody titers against the Beta and Gamma variants compared to the W+W group.

We also explored the possibility of using the Beta variant version of S-2P adjuvanted with CpG 1018 and aluminum hydroxide as the third dose in Group E (W + W + B). As shown in Figure 2, at five weeks after the third injection of Group E hamsters, NT_50_ GMTs were 6643, 1889, 1034, 1306, and 3595 against the wildtype, Alpha, Beta, Gamma, and Delta variants, respectively. Compared to the neutralizing titer against the wildtype, that against the Alpha, Beta, Gamma, and Delta variants had a 3.52-, 6.42-, 5.09- and 1.85-fold reduction, respectively. Two doses of wildtype S-2P combined with CpG and aluminum hydroxide followed by third dose of the adjuvanted Beta variant S-2P induced the best neutralization effect against the wildtype and all of the VoCs tested when compared to the other groups, especially against the Delta variant. The dosing regimens in Groups A to E resulted in a 1.9-3.8-fold lower NT_50_ GMT against the Delta variant than against the wildtype; however, the NT_50_ titers against the wildtype were different in each group. Unexpectedly, the Beta variant of S-2P appears to be the most suitable candidate for the third dose booster.

Pseudovirus neutralization assays were performed with lentivrus pseudo-typed with spike protein from the wildtype, Alpha, Beta, and Delta variants. Similar to the results of live neutralization assay, the highest levels of neutralizing antibody titers against the wildtype and variants were all found in the group receiving two doses of wildtype S-2P followed by one dose of the Beta variant S-2P (Group E - Figure 3). All groups immunized with S-2P produced high levels of neutralizing antibody against pseudoviruses of wildtype, Alpha, and Beta variants (Groups A to E). However, only hamsters receiving two doses of wildtype S-2P followed by one dose of Beta variant S-2P (Group E) produced high levels of neutralizing antibody against Beta and Delta andvariant pseudoviruses. In contrast, hamsters receiving two doses of S-2P (Groups A to C) had lower neutralizing antibody levels against the Beta and Delta variant pseudoviruses, whereas Group B produced the lowest level of antibodies against the Delta variant pseudovirus (Figure 3). The results of pseudovirus neutralization assays were similar to the live virus neutralization assays in that the administration of two doses of wildtype S-2P followed by one dose of Beta variant S-2P provided the broadest spectrum of immunity against wild-type SARS-CoV-2 and VoCs. After the serum samples have been tested against Alpha, Beta, and Delta variant pseudoviruses, the remaining samples from Groups D and E were tested against the Omicron variant pseudovirus. In Figure 4, although the neutralization titers against Omicron were reduced dramatically in both groups, Group E (6.8-fold) showed less reduction than that of Group D (17.8-fold). Boosting with the Beta variant S-2P was able to increase ID_50_ GMT against Omicron by 3.8 times and 1.5 times against the wildtype (Figure 4). Taken together, two doses of wildtype S-2P followed by a booster dose of Beta variant S-2P could increase immunity against VoCs, including the Omicron variant, compared to three doses of wildtype S-2P.

**Figure 3.**
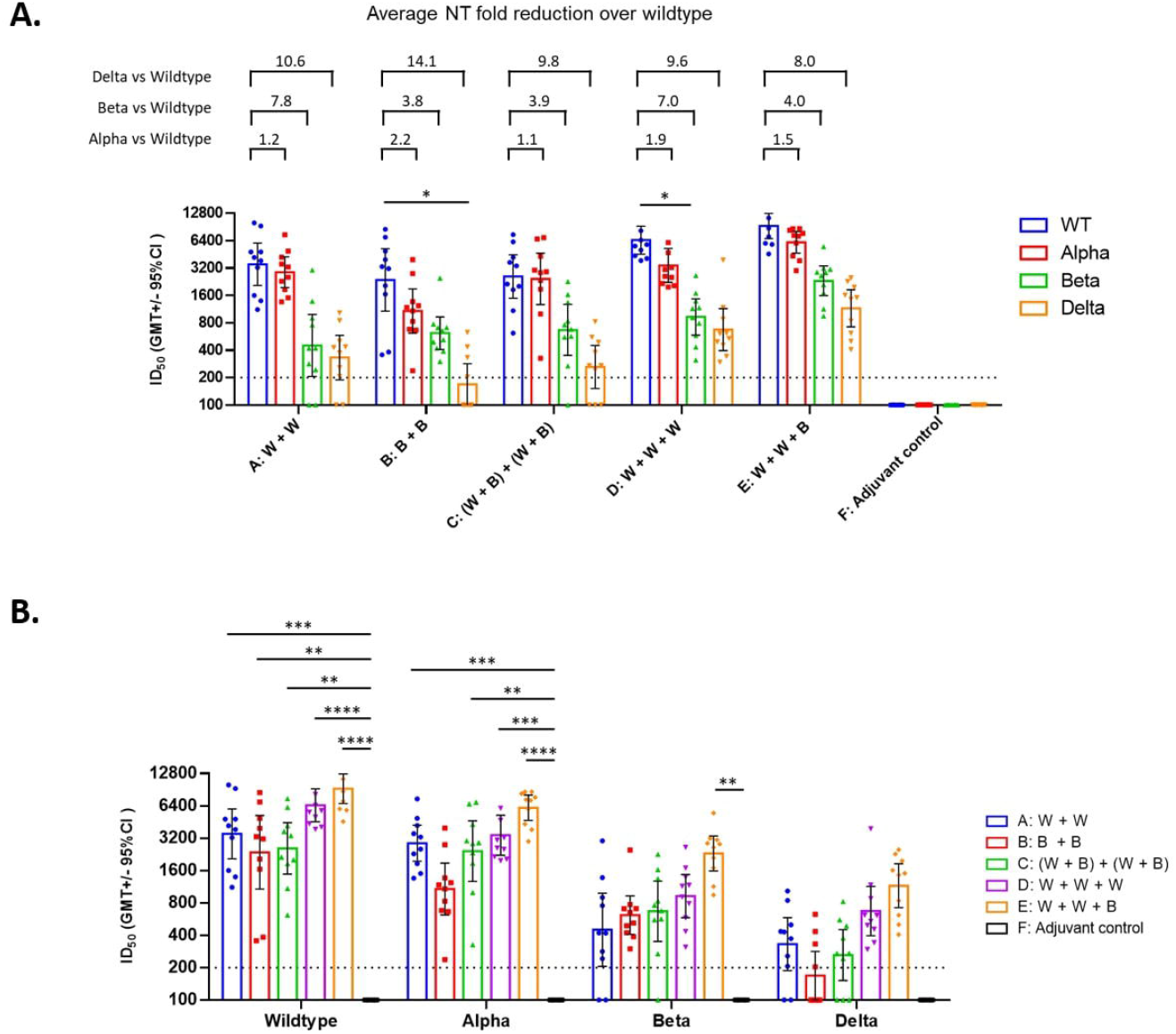
Neutralizing antibody titers with pseudovirus neutralization assay in hamsters five weeks after the final immunization. Hamsters were immunized and serum samples taken as in Figure 2. The samples were tested against lentivirus pseudotyped with the spike proteins of SARS-CoV-2 wildtype, Alpha, Beta, Delta, and Omicron variants. The results were **A:** Grouped according to vaccine regimen; **B:** Grouped according to variants. Bars indicate ID_50_ GMT with individual values displayed as symbols and error bars showing the 95% confidence intervals. Average fold reduction in GMT of variants against the wildtype strain were calculated and shown above brackets above the corresponding bars. Dotted line indicate the starting dilution (200) and all values below 200 are tabulated as 100. Statistical significance was calculated with Kruskal-Wallis with corrected Dunn’s multiple comparisons test. * = p < 0.05, ** = p < 0.01, *** = p < 0.001, **** = p < 0.0001

**Figure 4.**
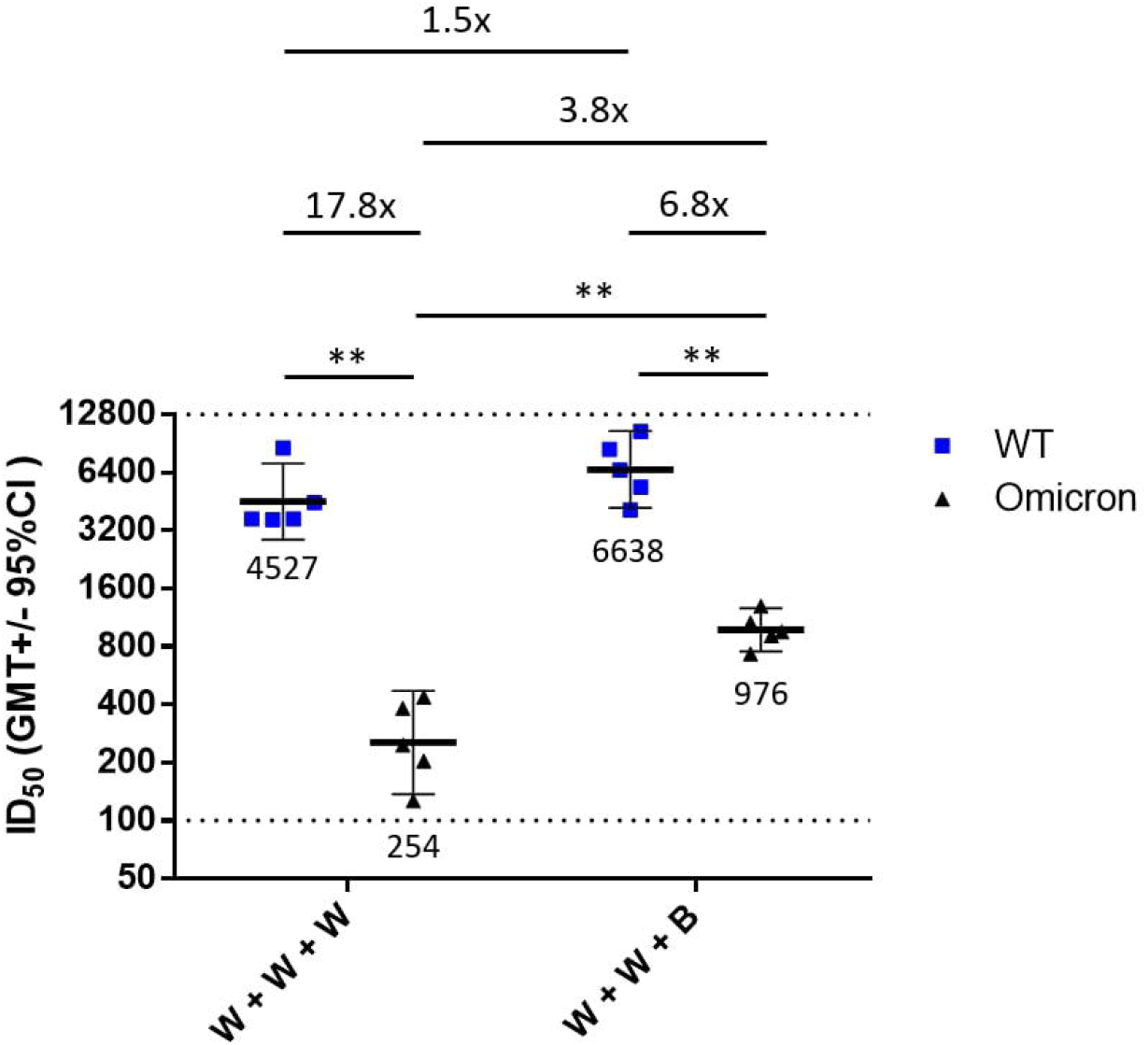
Neutralizing antibody titers with wildtype and Omicron variant pseudovirus neutralization assay. The serum samples from two hamsters were pooled together to form a sample size of n = 5 for each group. The pooled samples were tested against wildtype and Omicron variant pseudovirus neutralization assay. Horizontal bars indicate the ID_50_ GMT with individual ID_50_ values displayed as symbols and error bars showing the 95% confidence intervals. Fold difference in GMT comparisons were shown above brackets above the corresponding bars. Dotted line indicate the starting dilution (100) and the final dilution (12800) for the assay. Statistical significance was calculated with Mann-Whitney U test. * = p < 0.05, ** = p < 0.01, *** = p < 0.001, **** = p < 0.0001

### Protection from the Delta variant challenge in hamsters immunized with two doses of MVC-COV1901 or combined with a third dose of MVC-COV1901 or its Beta variant version of S-2P

Eight weeks after completion of the last immunization, hamsters were challenged with 10^4^ PFU of the Delta variant and body weights were monitored up to six days post infection (d.p.i.). All the vaccinated groups showed no significant weight loss six days after virus challenge in contrast to the adjuvant control group that showed a steady decline in weight during this period (Figure 5). The protection in all vaccinated hamster groups was significant (p < 0.0001) at 6 d.p.i. as compared to the weight loss seen in the adjuvant control group.

**Figure 5.**
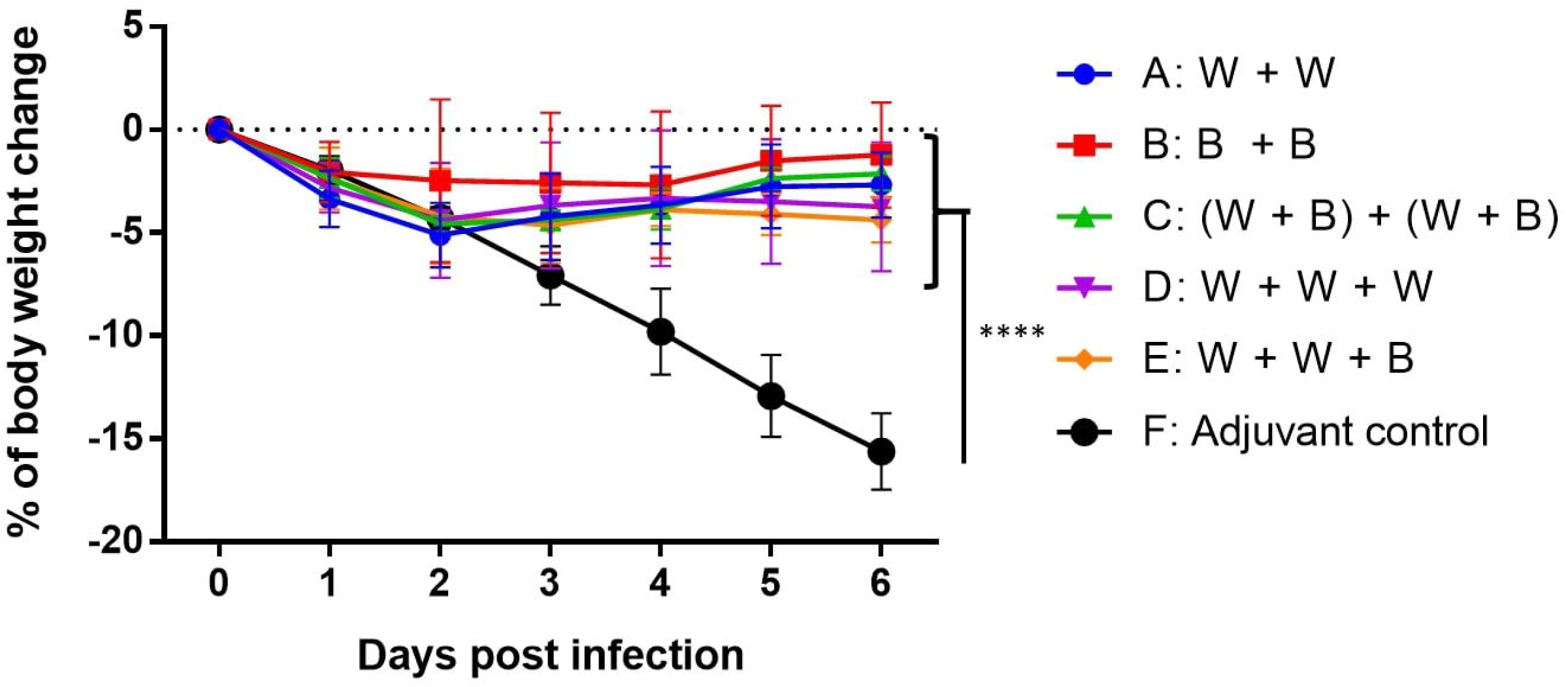
Change in body weight in hamsters after infection with SARS-CoV-2 Delta variant. Hamsters were challenged with SARS-CoV-2 Delta variant eight weeks after the final immunization. The body weights of individual hamsters were tracked daily up to the time of euthanizing at six days post infection. (n = 5/group). Results are shown as percent of weight relative to the day of challenge (day 0). Statistical significance was calculated with two-way ANOVA with Dunnett multiple comparison test with adjuvant only as a control. * = p < 0.05, ** = p < 0.01, *** = p < 0.001, **** = p < 0.0001

Lung viral load was measured by viral RNA and 50% tissue culture infectious dose (TCID_50_) assays. Figure 6A shows that at 3 d.p.i., lung viral RNA in Groups A to E hamsters were lower than in that of the adjuvant control group, but only in Group E was the lung viral RNA significantly lower than that in the adjuvant control group (p < 0.01), while by 6 d.p.i., the viral RNA in all groups were significantly (p < 0.05) lower than that of the control. In contrast, the viral titers in all of the vaccinated hamsters measured by TCID_50_ were significantly lower (p < 0.05) than that of the adjuvant control group at 3 d.p.i. (Figure 6B). Note that the lung viral load in hamsters, both viral RNA and especially viral titer as measured by TCID_50_, dropped considerably at 6 d.p.i. in the adjuvant-only control group likely due to hamsters’ natural immune response against the virus (Figure 6B). We also found a strong negative correlation (Spearman *r_s_* = −0.8227) between NT_50_ titer against the Delta variant from serum sampled five weeks after the final immunization and the number of viral genome at 3 d.p.i. (Figure S1). To examine the lung histopathology of the hamsters, lung necropsy sections were analyzed, and pathology scoring was tabulated at 3 d.p.i. or 6 d.p.i. (Figures 7, S2). There were no differences at 3 d.p.i. between control and experimental groups; however, at 6 d.p.i., the adjuvant control group had significantly (p < 0.01) increased lung pathology including extensive and severe immune cell infiltration, hemorrhage, and diffuse alveolar damage, compared to groups receiving three doses of S-2P (Groups D and E).

**Figure 6.**
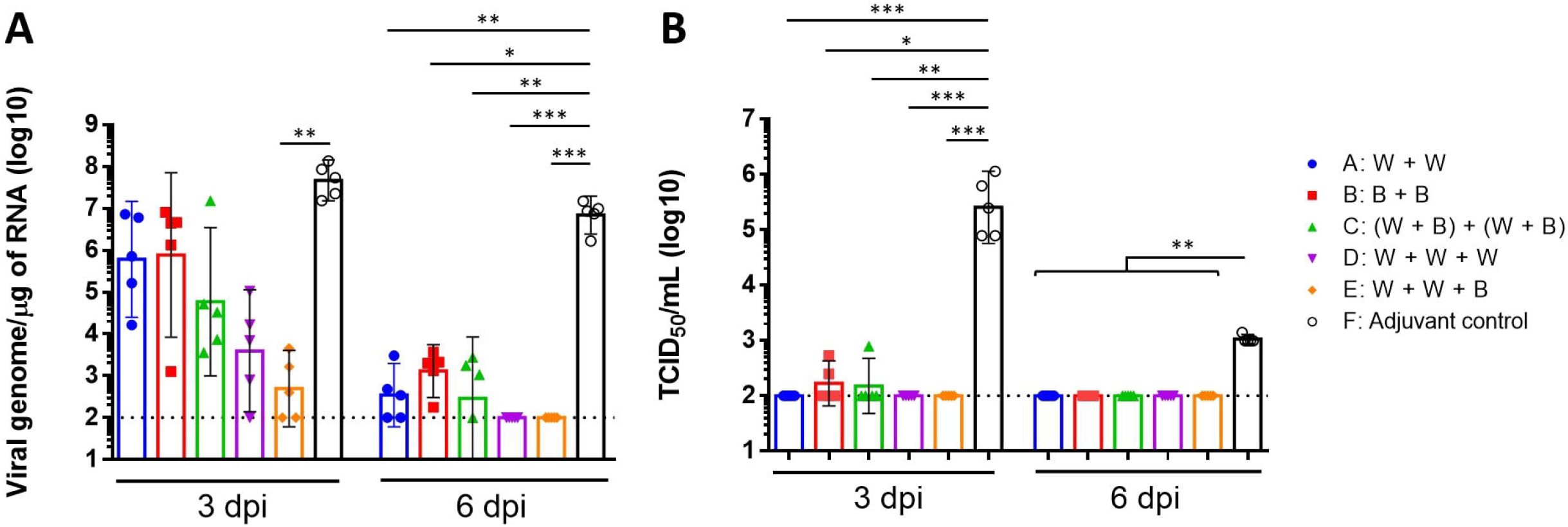
Viral load in hamsters three or six days post infection with SARS-CoV-2 Delta variant. The hamsters were euthanized at three or six days (n = 5/group) after infection and lung tissue samples were collected for viral load determination by “**A**” quantitative PCR of viral genome RNA, and “**B**” TCID_50_ assay for virus titer. Results are presented as geometric mean values with error bars representing 95% confidence intervals. Dotted line indicate limit of detection (200) and all bellows value the limit detection are tabulated as 200. Statistical significance was calculated with Kruskal-Wallis corrected Dunn’s multiple comparisons test. * = p < 0.05, ** = p < 0.01, *** = p < 0.001, **** = p < 0.0001

**Figure 7.**
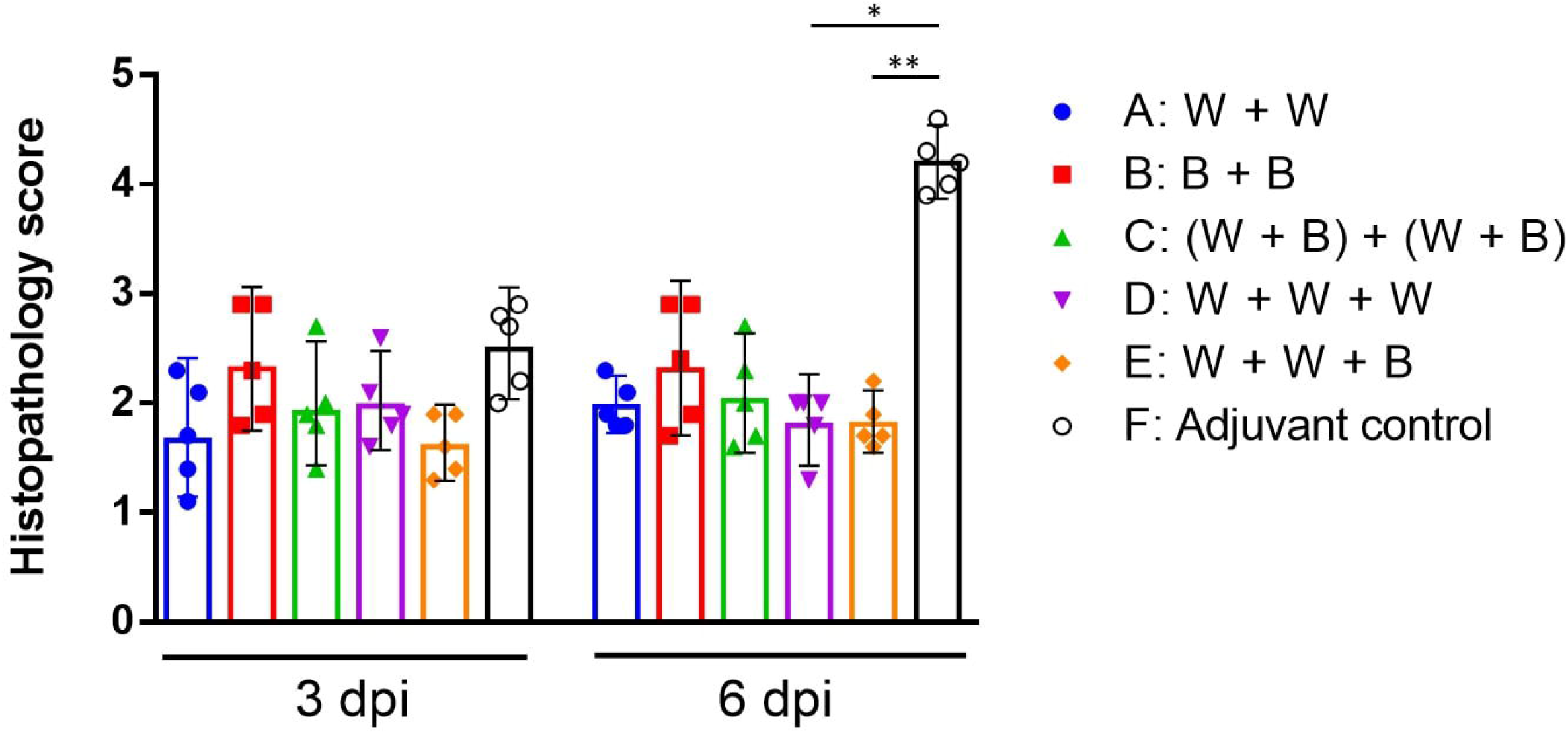
Lung histopathology scoring in hamsters at 3 or 6 days post infection with SARS-CoV-2 Delta variant. The hamsters were challenged with SARS-CoV-2 Delta variant and euthanized at three or six days after infection as in Figure 5. Lung sections were prepared and stained and histopathology scores were calculated. Results are presented as mean with error bars representing standard deviation. Statistical significance was calculated with Kruskal-Wallis corrected Dunn’s multiple comparisons test. * = p < 0.05, ** = p < 0.01, *** = p < 0.001, **** = p < 0.0001

## Discussion

We have previously shown that vaccination with both low and high dose of S-2P were effective against live SARS-CoV-2 virus challenge [16]. In the current study we extended our concept to examine the effect of a variant-based booster vaccinations followed by challenge with the Delta variant. Since Delta has emerged as the globally dominant variant at the time of hamster challenge before the appearance of the Omicron variant, we have chosen it as our model virus for challenge infection [21]. The results of the immunogenicity studies showed that immunization with either two doses of wildtype S-2P or two doses of the Beta variant S-2P induced lower level of antibody response against several of the VoCs tested. Wildtype S-2P fared poorly against the Beta and Gamma variants, whereas Beta variant S-2P induced higher neutralizing titers against only the Beta variant (Figure 2). A bivalent mixture of both wildtype and Beta variant S-2Ps showed results similar to immunization with two doses of wildtype S-2P, but with slightly increased immunogenicity against the Beta and Gamma variants. Three doses of wildtype S-2P was able to boost the titers against both Beta and Gamma more than the bivalent vaccine and also increased the neutralization titers against the Alpha and Delta variants as well. Unexpectedly, our study showed that the use of Beta variant S-2P as a third dose booster induced the highest and broadest spectrum of neutralizing titers against all variants as well as the wildtype. The recent surge of cases in fully vaccinated individuals due to the Omicron variant has warranted the use of boosters to address the waning immunity after vaccination and resistance of Omicron variant against antibody-mediated immunity [1, 8–10, 22]. Two doses of currently available vaccines performed poorly against the Omicron variant with many samples unable to neutralize the variant, but addition of a booster dose was able to improve the titer levels [9–10]. In this study we report that two doses of wildtype S-2P followed by a dose of Beta variant S-2P significantly increased neutralizing antibody titer against the Omicron variant pseudovirus compared to three doses of wildtype S-2P (Figure 4). Once the live Omicron variant virus is available, we intend to conduct a more throughout study on immunogenicity against the Omicron variant in hamsters.

All five regimens of vaccination protected hamsters from weight loss induced by infection with the Delta variant (Figure 5). Notably, while group B had a relatively poor antibody response against the Delta variant, this group did not experience any weight loss or increase in lung pathology (Figures 2, 5, and 7). In addition, the viral titers for the Delta variant in Group B were significantly lower than those of the adjuvant control, suggesting that the amount of anti-Delta antibodies and/or T cell immune responses induced by two doses of the Beta variant S-2P could have reduced viral replication in the lungs and protected the hamsters from weight loss and lung pathology (Figures 5 and 7). Since neutralizing antibodies induced by vaccines are polyclonal, they could also be cross-reactive with shared epitopes between different variants. Polyclonal antibodies induced by SARS-CoV-2 spike mRNA vaccine were profiled and were found to consist of a mixture of antibodies targeting the N-terminal domain (NTD) and the RBD, and they differ in their binding and neutralizing abilities [23, 24]. The re-stimulation of immunity may explain the low viral RNA titer in hamsters immunized in our study with either of the three-dose regimens. Further, the TCID_50_ live virus titers in all groups were very low and almost undetectable in most instances (Figure 6). This may be due to the sensitivity of the TCID_50_ assay itself, or the viral RNA assay may be detecting fragments of viral RNA from dead viruses as opposed to live replicating viruses. In future studies, subgenomic RNA detection should also be used to detect replicating viruses to corroborate the TCID_50_ results. The establishment of correlates of protection using the relationship between NT_50_ titer and viral RNA in a given hamster challenge model will help facilitate the expedited evaluation of vaccine combinations in future studies. Comparing neutralization titers against different viruses is complicated, as the assays used are inherently dissimilar. Nevertheless, our immunogenicity results demonstrated a clear correlation between neutralization titers and lung viral clearance in the hamsters (Figure S1).

Here we may propose a mechanism of which the administration of a different version of antigen as a booster dose would offer increased immunogenicity as compared to boosting with identical antigen. As illustrated in Figure S3, the spike protein is composed of S1 (head) and S2 (stem) domains. S1 contains the highly variable receptor binding motif (RBM) and is strain specific and immunodominant; while S2 is immunorecessive due to shielding by S1 but contain epitopes which are conserved across coronaviruses [25]. Other conserved regions in S1 such as the residues conserved in the RBD and N-terminus domain could also be targeted by broadly neutralizing antibodies [26]. The first two doses of wildtype S-2P would induce variant-specific antibodies to the S1 domain and broadly reactive antibodies against the S2 and other conserved domains. A third dose of wildtype S-2P would further induce more anti-wildtype S1 antibodies due to clonal expansion of anti-S1 memory B cells. Whereas a third dose consisting of Beta variant S-2P would allow the B cell response to select for antibodies that are reactive against epitopes common to the wildtype and Beta variant in the S2 and other conserved regions resulting in production of high affinity broadly neutralizing antibodies due to antibody affinity maturation. This immune refocusing has been studies in influenza hemagglutinin (HA) in which the glycosylation sites on the HA head has been removed to “unmask” the HA stem and to allow refocusing of B cell response to produce broadly neutralizing antibodies [27]. Further studies, e.g., profiling epitope-specific antibodies elicited by our vaccine, need to be done in order to support the potential mechanism we proposed here.

One limitation of this study is that we have not tested the vaccine’s protection in vivo with other VoCs besides the Delta variant; the vaccine efficacy against other VoCs is inferred from the neutralizing antibody titers. The natural course of infection among the hamsters includes a convalescent state, so the model does not allow for evaluating mortality or severe disease as endpoints. The lung histopathology scoring system we used in our animal model did not allow to distinguish different levels of lung damage caused by different degree of viral replication in the lung. T-cell functions were also not evaluated in the hamsters in this study, limiting our ability to assess the role of cellular immunity in providing protection. Two recent studies investigated the effects of a booster dose of ChAdOx1 and mRNA-1273 [13, 28]. Administration of a third dose using the Beta variant version of mRNA-1273 (mRNA-1273.351) following two doses of mRNA-1273 increased immunogenicity against the Beta variant more than did three doses of mRNA-1273. The administration of either mRNA-1273 or mRNA1273-351 as third dose exponentially boosted immunogenicity against Beta, Gamma, and Delta variants tested compared to two doses of mRNA1273 [13]. Concerning similar studies with ChAdOx1 vaccine, the third dose boosted neutralization titers against the Beta and Delta variants as well as gamma-interferon levels [28]. These findings are similar to our results that showed a third dose of vaccination could boost an immune response against the virus as well as its variants. Our study also showed that a 3^rd^ booster dose with homologous but especially with a heterologous Beta variant S-2P, increased immunogenicity against all the VoCs tested. Extrapolation from the unexpected and surprising hamster results to humans need to be supported by clinical evidence, for which the preparation is in progress. More recent data also support the use of boosting with vaccines such as mRNA1273, BNT162b2, and NVX-CoV2373 to generate anti-Omicron neutralizing response where two doses of vaccinations failed to neutralize the Omicron variant [6, 10, 29]. Findings from this study provide evidence to support the further evaluation of both the original and a Beta variant S-2P vaccine as a booster dose for individuals fully vaccinated with MVC-COV1901 as well as other approved vaccines.

## Acknowledgements

We would like to thank the team members at TFBS Bioscience Incorporation for their assistance with hamster housing and the immunization process. We also thank Academia Sinica for the use of their Biosafety Level 3 Facility and for providing the environment needed for handling and performing the SARS-CoV-2 hamster challenge, live SARS-CoV-2 neutralization assays and RNAi core facility pseudovirus neutralization assays. We also thank Drs. Han van den Bosch, Alexander Waits, and Michael Kornacker for reviewing and providing helpful inputs with the manuscript.

## Author Contributions

T.-Y. K., C.-C. W., W.-H. T, and J.C. produced the wildtype and Beta variant versions of S-2P antigens and pseudoviruses used in the study. T.-Y. K., C.-E. L., Y.-J. L., M.-Y. L., C.-C. W, W.-H. T., Y.-S. C., and C. C. designed the study and experiments. Y.-J. L. and Y.-S. C. supervised the experiments at TFBS Bioscience and Academia Sinica. Y.-J. L., M.-Y.-L., Y.-S. C., and L. T.-C. L. analyzed the results. M.-Y. L., J. D. C., P. T., Y.-S. C., and L. T.-C. L. drafted the manuscript. All authors reviewed and approved of the final version of the manuscript.

---

**Figure S1.**
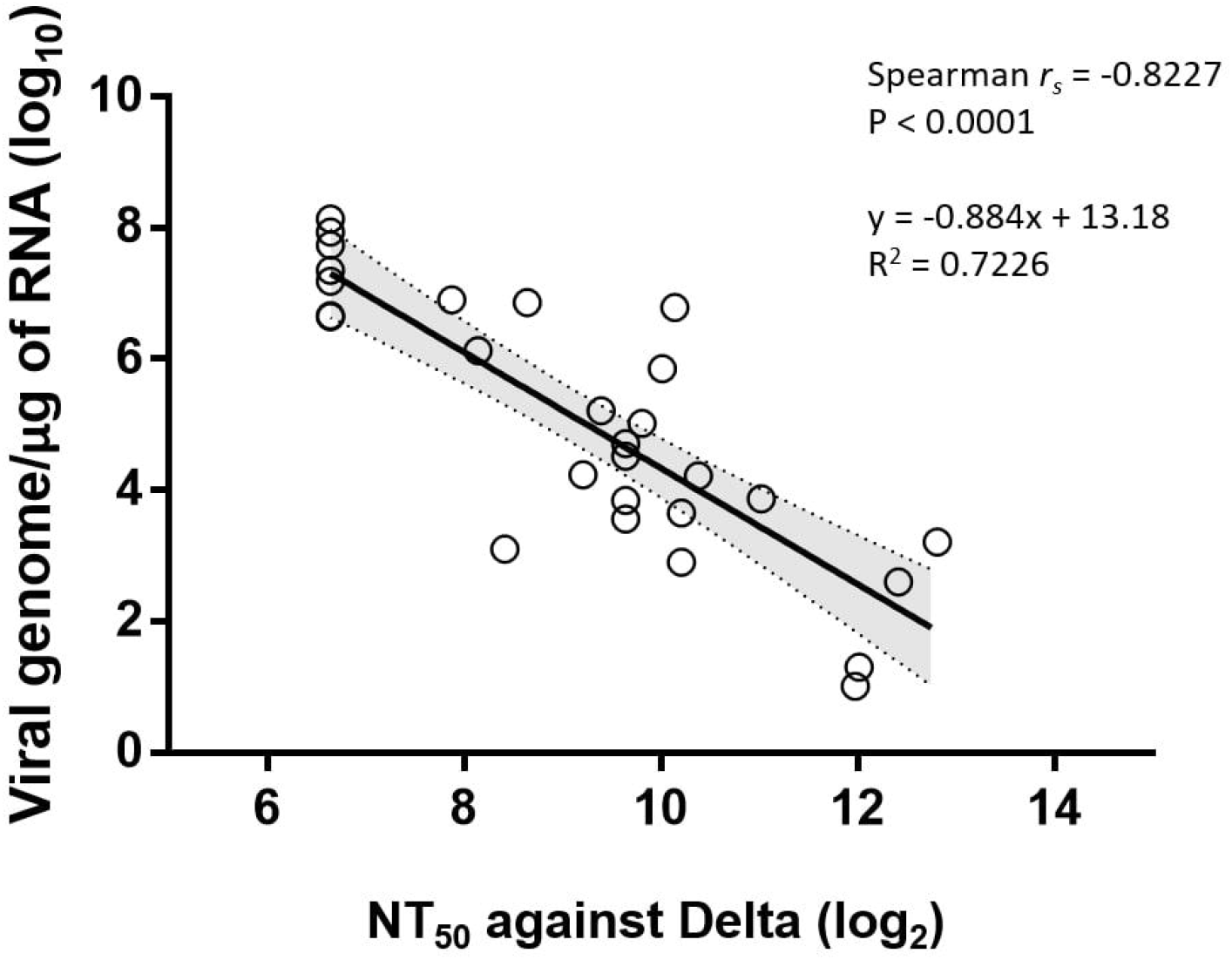
Correlation between SARS-CoV-2 viral genome copy numbers and NT_50_ titers against the Delta variant. Values of viral genome copy numbers 3 days post infection and NT50 titers against the Delta variant five days after the final immunization were tabulated (n = 29). Spearman’s rank correlation coefficient and linear regression were calculated with dotted bands and shaded area representing the 95% confidence bands of the linear regression line.

**Figure S2.**
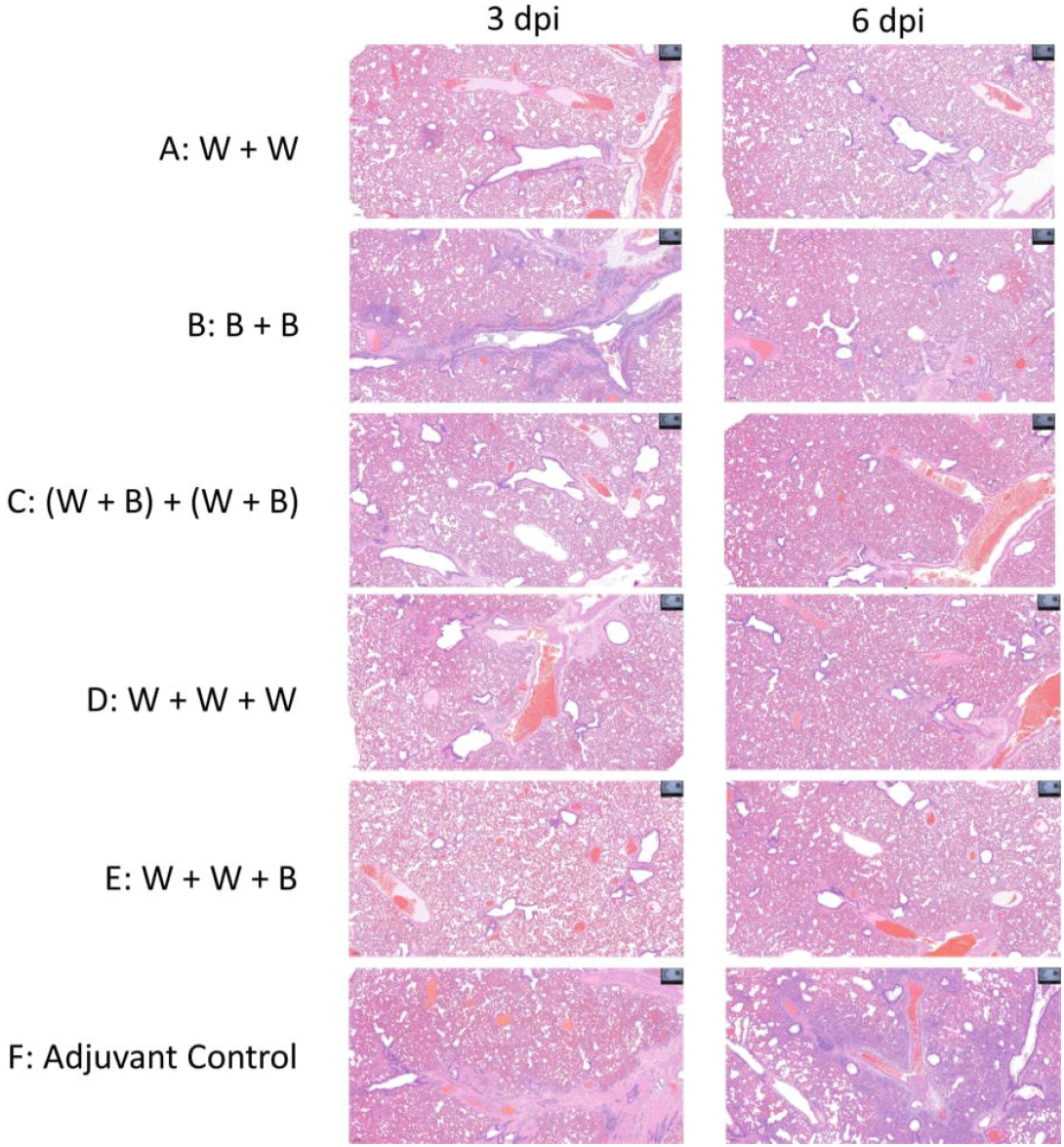
Representative histopathology sections in individual infected hamsters in the each of the vaccine groups at 3 d.p.i. or 6 d.p.i.

**Figure S3.**
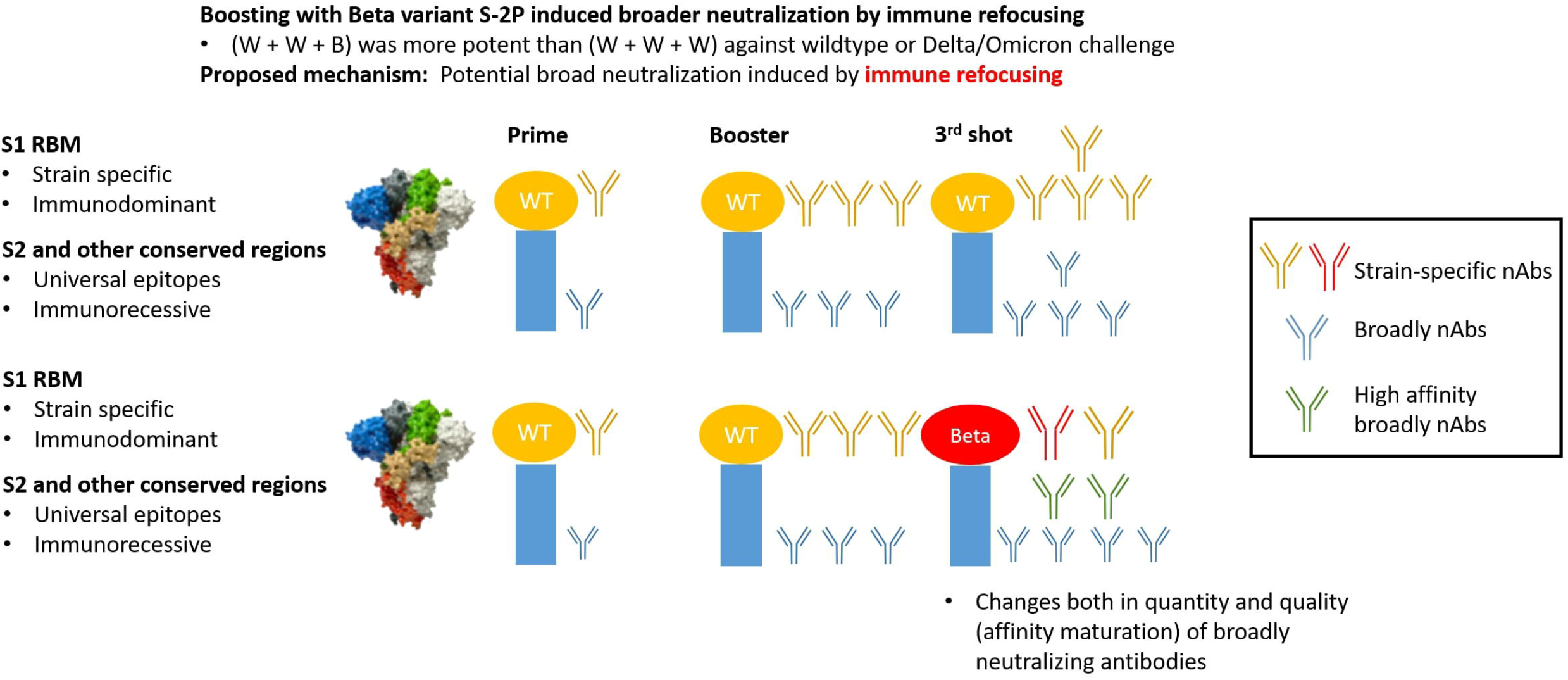
Proposed mechanism for immune refocusing.

## References

1. UK Health Security Agency. SARS-CoV-2 variants of concern and variants under investigation in England. Technical briefing: Update on hospitalization and vaccine effectiveness for Omicron VOC-21NOV-01 (B.1.1.529). December 31, 2021. Available at: https://assets.publishing.service.gov.uk/government/uploads/system/uploads/attachment_data/file/1044481/Technical-Briefing-31-Dec-2021-Omicron_severity_update.pdf. Retrieved January 3, 2022

2. World Health Organization. Tracking SARS-CoV-2 variants. Available at: https://www.who.int/en/activities/tracking-SARS-CoV-2-variants/. Accessed 15 September, 2021.

3. Choi A, Koch M, Wu K, et al. Serum neutralizing activity of mRNA-1273 against SARS-CoV-2 variants. bioRxiv. 2021 Jan 1.

4. Jongeneelen M, Kaszas K, Veldman D, et al. Ad26. COV2. S elicited neutralizing activity against Delta and other SARS-CoV-2 variants of concern. bioRxiv. 2021 Jan 1.

5. Liu C, Ginn HM, Dejnirattisai W, et al. Reduced neutralization of SARS-CoV-2 B. 1.617 by vaccine and convalescent serum. Cell. 2021 Aug 5;184(16):4220–36.

6. Carreno JM, Alshammary H, Tcheou J, Singh G, Raskin A, Kawabata H, Sominsky L, Clark J, Adelsberg DC, Bielak D, Gonzalez-Reiche AS. Activity of convalescent and vaccine serum against a B. 1.1. 529 variant SARS-CoV-2 isolate. medRxiv. 2021 Jan 1.

7. Emary KR, Golubchik T, Aley PK, et al. Efficacy of ChAdOx1 nCoV-19 (AZD1222) vaccine against SARS-CoV-2 variant of concern 202012/01 (B. 1.1. 7): an exploratory analysis of a randomized controlled trial. The Lancet. 2021 Apr 10;397(10282):1351–62.

8. Madhi SA, Baillie V, Cutland CL, et al. Efficacy of the ChAdOx1 nCoV-19 Covid-19 vaccine against the B. 1.351 variant. New England Journal of Medicine. 2021 May 20;384(20):1885–98.

9. Lu L, Mok BW, Chen L, Chan JM, Tsang OT, Lam BH, Chuang VW, Chu AW, Chan WM, Ip JD, Chan BP. Neutralization of SARS-CoV-2 Omicron variant by sera from BNT162b2 or Coronavac vaccine recipients. medRxiv. 2021 Jan 1

10. Planas D, Saunders N, Maes P, Benhassine FG, Planchais C, Porrot F, Staropoli I, Lemoine F, Pere H, Veyer D, Puech J. Considerable escape of SARS-CoV-2 variant Omicron to antibody neutralization. bioRxiv. 2021 Jan 1

11. Puranik A, Lenehan PJ, Silvert E, et al. Comparison of two highly-effective mRNA vaccines for COVID-19 during periods of Alpha and Delta variant prevalence. medRxiv. 2021 Aug 8.

12. Lopez Bernal J, Andrews N, Gower C, et al. Effectiveness of Covid-19 vaccines against the B. 1.617. 2 (delta) variant. New England Journal of Medicine. 2021 Jul 21.

13. Choi A, Koch M, Wu K, Chu L, Ma L, Hill A, Nunna N, Huang W, Oestreicher J, Colpitts T, Bennett H. Safety and immunogenicity of SARS-CoV-2 variant mRNA vaccine boosters in healthy adults: an interim analysis. Nature medicine. 2021 Nov;27(11):2025–31.

14. Mallory R, Formica N, Pfeiffer S, Wilkinson B, Marcheschi A, Albert G, McFall H, Robinson M, Plested J, Zhu M, Cloney-Clark S. Immunogenicity and Safety Following a Homologous Booster Dose of a SARS-CoV-2 recombinant spike protein vaccine (NVX-CoV2373): A Phase 2 Randomized Placebo-Controlled Trial. medRxiv. 2021 Jan 1.

15. Kuo TY, Lin MY, Coffman RL, et al. Development of CpG-adjuvanted stable prefusion SARS-CoV-2 spike antigen as a subunit vaccine against COVID-19. Scientific reports. 2020 Nov 18;10(1):1–0.

16. Lien CE, Lin YJ, Chen C, et al. CpG-adjuvanted stable prefusion SARS-CoV-2 spike protein protected hamsters from SARS-CoV-2 challenge. Scientific reports. 2021 Apr 22;11(1):1–7.

17. Hsieh SM, Liu WD, Huang YS et al. Safety and Immunogenicity of a Recombinant Stabilized Prefusion SARS-CoV-2 Spike Protein Vaccine (MVC-COV1901) Adjuvanted With CpG 1018 and Aluminum Hydroxide in Healthy Adults: A Phase 1, Dose-Escalation Study. EClinicalMedicine. June 26, 2021.

18. Szu-Min H, Liu MC, Chen YH, et al. Safety and Immunogenicity of CpG 1018 and Aluminum Hydroxide-Adjuvanted SARS-CoV-2 S-2P Protein Vaccine MVC-COV1901: A Large-Scale Double-Blind, Randomised, Placebo-Controlled Phase 2 Trial. medRxiv. 2021 Jan 1.

19. Blanchard B and Lee Y. “Taiwan approves Medigen’s COVID-19 vaccine candidate”. July 19, 2021. Available at: https://www.reuters.com/world/asia-pacific/taiwan-approves-production-medigens-covid-19-vaccine-candidate-2021-07-19/”. Accessed September 17, 2021.

20. Lien CE, Kuo TY, Lin YJ, et al. Evaluating the neutralizing ability of a CpG-adjuvanted S-2P subunit vaccine against SARS-CoV-2 Variants of Concern. Clin Infect Dis. 2021. In press.

21. Julia LM, Ginger T, Alaa AL, et al. Delta Variant Report, outbreak.info. Available online: https://outbreak.info/situation-reports/delta. Accessed September 17, 2021.

22. Levin EG, Lustig Y, Cohen C, Fluss R, Indenbaum V, Amit S, Doolman R, Asraf K, Mendelson E, Ziv A, Rubin C. Waning immune humoral response to BNT162b2 Covid-19 vaccine over 6 months. New England Journal of Medicine. 2021 Dec 9;385(24):e84.

23. Amanat F, Thapa M, Lei T, et al. The plasmablast response to SARS-CoV-2 mRNA vaccination is dominated by non-neutralizing antibodies that target both the NTD and the RBD. medRxiv. 2021 March 9.

24. Tong P, Gautam A, Windsor I, et al. Memory B cell repertoire for recognition of evolving SARS-CoV-2 spike. bioRxiv. 2021 March 10.

25. Huang Y, Nguyen AW, Hsieh CL, Silva R, Olaluwoye OS, Wilen R, Kaoud TS, Azouz LR, Qerqez AN, Le KC, DiVenere AM. Identification of a conserved neutralizing epitope present on spike proteins from highly pathogenic coronaviruses. bioRxiv. 2021 Jan 1.

26. Cameroni E, Saliba C, Bowen JE, Rosen LE, Culap K, Pinto D, De Marco A, Zepeda SK, di Iulio J, Zatta F, Kaiser H. Broadly neutralizing antibodies overcome SARS-CoV-2 Omicron antigenic shift. bioRxiv. 2021 Jan 1.

27. Liu WC, Jan JT, Huang YJ, Chen TH, Wu SC. Unmasking stem-specific neutralizing epitopes by abolishing N-linked glycosylation sites of influenza virus hemagglutinin proteins for vaccine design. Journal of virology. 2016 Sep 12;90(19):8496–508.

28. Flaxman A, Marchevsky NG, Jenkin D, et al. Reactogenicity and immunogenicity after a late second dose or a third dose of ChAdOx1 nCoV-19 in the UK: a substudy of two randomised controlled trials (COV001 and COV002). The Lancet. 2021 Sep 1.

29. Mallory R, Formica N, Pfeiffer S, Wilkinson B, Marcheschi A, Albert G, McFall H, Robinson M, Plested J, Zhu M, Cloney-Clark S. Immunogenicity and Safety Following a Homologous Booster Dose of a SARS-CoV-2 recombinant spike protein vaccine (NVX-CoV2373): A Phase 2 Randomized Placebo-Controlled Trial. medRxiv. 2021 Jan 1.

